# Matrix Metalloproteinase (MMP) inhibition rescues endothelial glycocalyx damage and reduces neutrophil infiltration in sepsis-associated acute kidney injury

**DOI:** 10.64898/2025.12.04.691889

**Authors:** Haije Wu, Laura Carey, Matthew J. Butler, Vaishnavi Lalam, Nesreen Hamad, Aldara Martin Alonso, Monica Gamez, Sevil Erarslan, Michael Crompton, Jasmine Aldam, Holly Stowell-Connolly, Gavin I. Welsh, Rebecca R. Foster, Simon C. Satchell, Raina D. Ramnath

## Abstract

Endothelial glycocalyx (eGlx), a carbohydrate-rich endothelial coat, maintains vascular homeostasis, and its disruption contributes to several vascular diseases. We previously identified matrix metalloproteinase (MMP) 2 and 9-mediated eGlx shedding as a key mechanism in human glomerular endothelial cell (GEnC) damage and kidney dysfunction in diabetes. Here, we sought to determine whether this mechanism contributes to renal and systemic microvascular endothelial cell injury in sepsis-AKI.

Sepsis-AKI was induced in mice by lipopolysaccharide (LPS) injection. Confocal microscopy demonstrated glomerular and peritubular eGlx damage, and increased circulating SDC4 demonstrated systemic eGlx shedding. These were associated with enhanced mRNA expression of eGlx components (syndecans 1 and 4) and endothelial inflammatory markers, Intercellular and Vascular Cell Adhesion Molecules (ICAM and VCAM) in the kidney. Additionally, sepsis-AKI markers, serum creatinine and urea, were increased. We found that renal and circulating MMP9, but not MMP2, levels were raised in sepsis-AKI. Moreover, MMP9 staining colocalises with neutrophil, endothelial and mesangial staining in glomeruli in sepsis-AKI. Treatment with MMP2 and 9 inhibitor 1, 1h before LPS injection, prevented glomerular, peritubular and systemic eGlx damage in sepsis-AKI. MMP inhibition also attenuated glomerular leucocyte numbers in sepsis-AKI. We confirmed increased MMP9 activity, shedding of eGlx syndecan 1 and elevated angiopoietin 2 levels, an endothelial damage marker, in sepsis-AKI in humans. Moreover, we used human glomerular endothelial cells (GEnC) in vitro to demonstrate that human sepsis-AKI serum directly damages GEnC eGlx.

Our studies confirm that MMP-mediated eGlx damage is a key contributor to renal and systemic microvascular dysfunction in sepsis-AKI and represents a potential therapeutic target for protecting the microvasculature.

## Introduction

Sepsis is a severe and dysregulated inflammatory host response to injury or infection. It is the leading cause of acute kidney injury (AKI), accounting for over 50% of patients with AKI in intensive care units.^1^ Patients with sepsis-AKI are more likely to require renal replacement therapy than patients with sepsis alone or with non-sepsis AKI.^2^ Sepsis-AKI is a major cause of death, both pre- and post-hospital discharge. Additionally, it significantly increases length of hospital stay, tripling healthcare costs compared with sepsis alone.^3^ Current treatments for sepsis-AKI remain supportive and nonspecific, with antibiotics and resuscitation fluids forming the cornerstones of therapy.^4^ Effective therapies targeting the pathobiology of sepsis-AKI are urgently required.

The vascular endothelium is a key player in the pathogenesis of sepsis-AKI. The kidney and systemic microcirculations, including glomerular and peritubular capillaries, are profoundly disturbed.^5^ This is associated with inflammatory cell infiltration, increased permeability, hemodynamic changes, and oedema, all of which contribute to kidney damage in sepsis.^1, 3, 5–7^ Endothelial glycocalyx (eGlx) is a hydrated poly-anionic gel, present on the luminal surface of all blood vessels, including in the renal microcirculation. It is essential for maintenance of vascular homeostasis and is damaged in disease states. The glycocalyx consists of proteoglycans, such as syndecan (SDC)1 and 4, glycosaminoglycans, such as heparan sulphate (HS), and glycoproteins, among other components.^8, 9^

Plasma SDC1 and HS levels are elevated in patients with sepsis, likely due to shedding of systemic eGlx.^10^ Their levels correlate with clinical severity scores and mortality.^10^ In addition, SDC4 has been identified as a novel biomarker that could predict sepsis severity and mortality in paediatric sepsis.^11^ EGlx shedding is associated with dysregulated endothelial cells in a human model of endotoxemia^12, 13^ and disrupted blood flow in a rodent model of sepsis,^5, 14^ demonstrating a clear association between eGlx damage and endothelial dysfunction in sepsis. Studies are emerging that show Glx shedding in urine and plasma is linked to kidney dysfunction and mortality in sepsis patients.^15, 16^ We have shown that glomerular eGlx constitutes a barrier to protein permeability and that its damage contributes to glomerular endothelial albumin permeability in vitro,^8, 17–19^ exvivo,^20–22^ and elevated urine albumin creatinine ratio in vivo ^23, 24^ Others have demonstrated increased leucocyte trafficking in the glomeruli^25–27^ and kidney dysfunction.^7^ Loss of eGlx on peritubular capillaries is associated with CKD.^9, 28^ In inflamed tissues, shedding of the eGlx exposes adhesion molecules and selectins to circulating leucocytes, promoting the process of transmigration.^29, 30^

Matrix metalloproteinases (MMP) act physiologically to cleave proteoglycans, as part of their normal turnover. Dysregulation of MMP2 and 9 activity is implicated in kidney,^31^ brain^32^ and liver^33^ damage in ischaemia-reperfusion injury models. Elevated circulating levels of MMP9 are reported in sepsis patients.^34^ However, the adverse effects of MMPs on the renal microcirculation in sepsis-AKI have not been explored in detail. We identified MMP2 and 9-mediated SDC4 shedding as a key mechanism of eGlx dysfunction in glomerular endothelial cells (GEnC), in response to the inflammatory cytokine, TNFα.^8^ Similarly, this mechanism was implicated in a mouse model of early diabetic kidney disease and in a Renin-Angiotensin-Aldosterone System (RAAS) model of kidney injury.^22, 35^ Here, we aim to investigate whether this mechanism contributes to renal and systemic microvascular endothelial cell injury, and whether inhibiting this process could protect against renal and systemic microvascular dysfunction, in sepsis-AKI. We therefore hypothesise that MMP2 and 9-mediated eGlx loss contributes to glomerular, peritubular and systemic microvascular dysfunction in sepsis-AKI and represents a potential therapeutic target for repairing microvascular damage in sepsis-AKI.

## Methods

All animal experiments were approved by the UK Home Office and carried out in accordance with established International Guiding Principles for Animal Research. C56BL6 mice were maintained in the Animal Housing Unit at the University of Bristol in an environment with controlled temperature (21–24°C) and lighting (12:12-h light–darkness cycle). Standard laboratory chow and drinking water were provided ad libitum. A period of 1-2 weeks was allowed for animals to acclimatise before any experimental manipulations were undertaken.

### Models

#### Endotoxin model

The endotoxemia model was induced by giving 10 mg/kg LPS or vehicle (saline) intraperitoneally to 8-week-old C57BL6 mice and they were culled at 18h post-LPS injections. In another cohort, 1 hour before LPS injection, mice were given a single intraperitoneal injection of 5mg/kg MMP2 and 9 inhibitor I (MMPI), biphenylylsulfonylamino-3-phenylpropionic acid^35–38^ (Merck, Middlesex, UK) or vehicle (0.05% DMSO in PBS) and the mice were culled at 18h after LPS injections. MMPI is a potent and highly selective inhibitor that binds to the zinc ion at the active site of gelatinases (MMP2 and 9), thereby blocking their activities.^39^ The inhibitor is potent at blocking MMP9 (IC50: 240nM) and MMP 2 (IC50: 310nM). Systolic and diastolic blood pressure (BP), were measured using a noninvasive tail cuff method, as previously.^35^ For BP measurement, 15 inflation cycles were used, and for inclusion, the computer had to accept 6 of the 15 readings. For tissue collection, mice were anaesthetised with 3.5% isoflurane in 1L/min O_2_. A midline laparotomy was performed, followed by blood collection directly from the left ventricle for serum/plasma separation. Cardiac perfusion with Ringer’s solution (NaCl, 132mM; KCl, 4.6mM; MgSO4-7H2O 1.27mM; CaCl2-2H2O 2mM; NaHCO3, 25mM; D(+)glucose, 5.5mM; N-2-hydroxyethylpiperazine-N’-2-ethanesulphonic (HEPES) acid, 3.07mM; HEPES sodium salt, 1.9mM, pH 7.40) was performed to flush both kidneys. One kidney was removed for glomerular sieving, as previously,^40^ and portions of the other kidney were fixed in 4% paraformaldehyde and snap-frozen for histology, ELISA and real-time PCR.

#### Study population

Sixteen adult sepsis-AKI patients were recruited from Southmead Hospital, Bristol, UK. Eighteen adult volunteers, recruited from hospital staff, served as healthy controls. All participants gave informed consent. Demographic and physiological variables are outlined in (**Table 1**). Blood samples from patients and controls were centrifuged at 1200 rcf for 10 mins at 4 °C. Serum was aliquoted and stored at – 80 °C to analyse SDC1, endothelial angiopoietin 2 (ANGPT2), and MMP9 activity.

**Table 1:** Baseline Characteristics for Controls and Patients.

| Table 1: Baseline Characteristics for Controls and Patients |  |  |  |  |
| --- | --- | --- | --- | --- |
| Variable | Category | Control | Sepsis | p-value |
| Number of participants |  | 18 | 16 |  |
| Age (years) |  | 58.80 (18.34) | 56.40 (18.21) | >0.05 |
| Sex | Male | 7 (38.9%) | 10 (62.5%) | >0.05 |
| BMI (kg/m2) |  | — | 29.02 (8.90) |  |
| Ethnicity | White |  | 14 (87.5%) |  |
| Smoking Status | Never |  | 7 (46.7%) |  |
| SOFA Score (last 24h) |  |  | 8.53 (3.29) |  |
| Lowest MAP (mmHg) |  |  | 60.17 (15.20) |  |
| Outcome | Died |  | 3 (18.8%) |  |
| Clinical variables |  |  |  |  |
| Serum Creatinine at Recruitment (μmol/L) |  | 75.17 (13.17) | 173.50 (87.97) | <0.0001 |
| eGFR (mL/min/1.73m²) |  | 83.11 (10.66) | 49.44 (27.86) | 0.0016 |
| uACR (mg/mmol) |  | 2.41 (2.08) | 23.57 (25.98) | <0.0001 |
| uPCR (mg/mmol) |  | 11.09 (7.91) | 181.44 (150.82) | <0.0001 |
| Lactate (mmol/L) |  | — | 9.67 (20.69) |  |
| Latest WCC (×10<sup>9</sup>/L) |  |  | 13.60 (6.77) |  |
| Highest WCC (×10<sup>9</sup>/L) |  |  | 25.55 (12.36) |  |
| Highest CRP (mg/L) |  |  | 287.94 (135.48) |  |
| Table 1: Numeric values are presented as mean (SD); categorical values are presented as n (%). Statistical tests used: Unpaired t-test was used on normally distributed data and if normality was not achieved, Mann-Whitney test was used. |  |  |  |  |

#### Definitions

Sepsis was defined according to Sepsis-3 as an acute increase of ≥2 in the Sequential Organ Failure Assessment (SOFA) score (**Table 1**) in patients with either documented or suspected infection.^41^ AKI and its severity were defined according to KDIGO 2012 guidelines:^42^ Sepsis-AKI patients met the criteria for inclusion for both sepsis and AKI (supplementary methods).

#### Sepsis-AKI in vitro model

Human GEnC were pretreated with Click-iT™ ManNAz (tetraacetylated N-Azidoacetyl-D-Mannosamine), to label newly synthesised eGlx,^43^ followed by stimulation with 10% serum from control participants or sepsis-AKI patients. EGlx was imaged and quantified (blind) using confocal microscopy and ImageJ software.

#### Lectin-based imaging of eGlx in mouse kidney sections

Five µm paraffin-embedded kidney sections were dewaxed in histoclear, followed by rehydration in graded ethanol (100%, 90% and 70%) and a wash in PBS. The sections were incubated in blocking buffer [1% BSA in PBS containing 0.5% Tween (PBX)] for 30 min. After 3 washes in PBX, endogenous biotin was blocked using a streptavidin/biotin blocking kit (SP-2002, Vector Laboratories) as per the manufacturer’s instructions. After 2 washes, the sections were incubated with biotinylated Lycopersicon esculentum lectin (LEL) (2mg/ml) 1:100 in blocking buffer, pH 6.8 overnight at 4°C. Buffer only was used as a negative control. After 4 washes, the sections were incubated with streptavidin AF488 (S32354, ThermoFisher Scientific, 1:500) in the blocking buffer, pH 6.8, 1 hour at room temperature. After 4 washes in PBS, the nuclei were counterstained with DAPI (Invitrogen; Life Technologies). The sections were incubated with an endothelial cell membrane label R18 (O246, ThermoFisher Scientific, at 1:1000 dilution) for 10 min. After a 50-second dip in PBS, the coverslips were mounted in Vectashield mounting medium (Vector Laboratories) and examined using Leica SP8 Acousto-Optical Beam Splitter confocal laser scanning microscope attached to a Leica DM I8 inverted epifluorescence microscope. Confocal images of kidney sections were taken using a 63x/1.40 oil objective with a zoom factor of 2.6 and format set to 1024×1024 to give pixel size of 69.38nm

#### EGlx depth: peak to peak analysis

EGlx depth was assessed using confocal images obtained as above and as previously described.^22, 44^ Briefly, a line was drawn from the inside to the outside of the glomerular and peritubular capillary loop, crossing the glycocalyx first, followed by the R18 endothelial membrane (Fig 1). The line is drawn perpendicular to the eGlx and the cell membrane label to get the maximum consistent depth of the glycocalyx. Fluorescence intensity profiles were then generated for both LEL, denoting eGlx, and R18, labelling endothelial membrane. The distance between the peak profile signals (peak to peak: P-P) is an index of glycocalyx depth. The same P-P quantification methodology was applied to quantify glomerular and peritubular eGlx depth. Glomerular eGlx depth was quantified using 5 glomeruli per mouse, with 3 capillary loops analysed per glomerulus and 4 measurement lines per capillary loop. Peritubular eGlx depth was assessed from 5 peritubular capillaries per mouse, with 4 lines measured per capillary loop.

**Figure 1.**
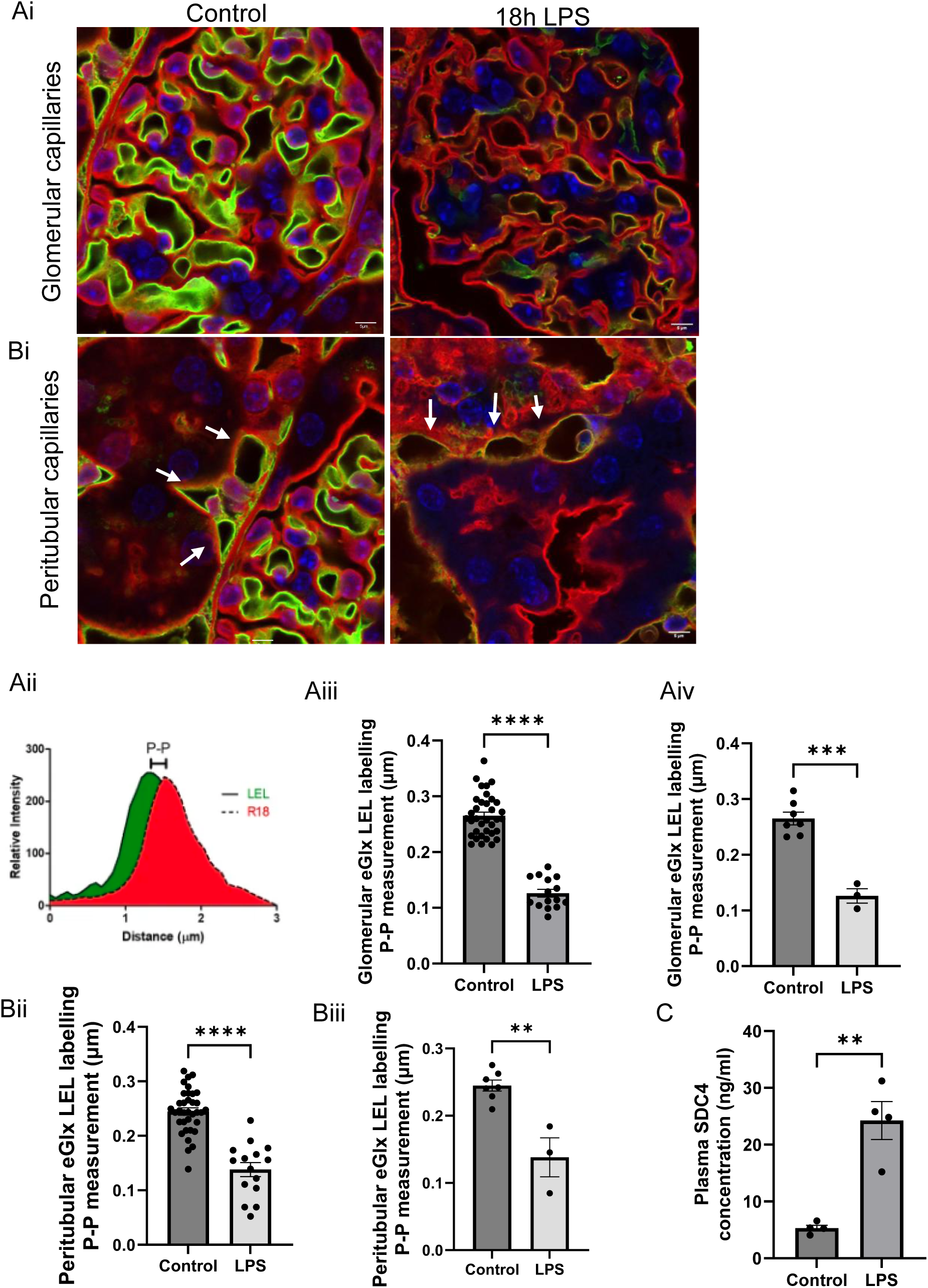
Glomerular, peritubular and systemic eGlx are damaged in sepsis-AKI. Representative image show (Ai) glomerular and (Bi) peritubular capillaries labelled red (R18) and the luminal eGlx labelled green (LEL). (Aii) The difference between the peaks (p-p) of the LEL and R18 fluorescence profiles is a measure of the eGlx depth. P-P assessment of eGLX depth in (Aiii, iv) glomerular capillary is control 0.2651 ± 0.01147 n=7 mice,18h LPS 0.1262 ± 0.01292 n=3 mice *** p=0.0001 (Bii, iii) peritubular capillary control 0.2448 ± 0.008255 n=7 mice, 18h LPS 0.1381 ± 0.02897 n=3 mice **p=0.0012. (C) Circulating SDC4 shedding in the plasma was determined at 18h post-LPS injections, control 5.319 ± 0.5056 n=4 mice, 18h LPS 24.25 ± 3.328 n=4 mice, **p=0.0014. Each dot on the graph represents a mouse. Data are expressed as the mean ± SEM. An unpaired t-test was used if the data were normally distributed. If normality was not achieved, Mann Whitney test was used for statistical analysis.

#### MMP9 immunofluorescence on mouse kidney sections

As MMP9, but not MMP2, level was elevated in sepsis-AKI in mice and humans, only MMP9 immunostaining was performed. Frozen mouse kidney sections (5 μm) were fixed in 4% PFA for 15 minutes and then blocked with 5% BSA for 1h at room temperature. The sections were incubated with goat anti-MMP9 antibody (1:100, R&D Systems, Minneapolis, USA) overnight at 4 °C. After 3 washes, the sections were incubated with donkey anti-goat secondary antibody (1:200, Thermo Fisher Scientific, MA, USA) for 1h at room temperature. MMP9-stained sections were co-stained with rabbit anti-Nephrin (1:100, Abcam, Cambridge, UK), rabbit anti-PDGFRβ (1:100, Cell Signalling Technology, Leiden, Netherlands), rat anti-endomucin (1:100, Santa Cruz, Texas, USA) and anti-calgranulin A (1:100, a gift from Dr Amulic,^45^ an in-house antibody developed at the Max Planck Institute for Infection Biology, Berlin) antibodies, for 1 hour at room temperature to identify podocyte, mesangial, endothelial cells and neutrophils, respectively. After 3 washes, sections were incubated in either donkey anti-rabbit or goat anti-rat secondary antibody (1:200, Thermo Fisher Scientific, MA, USA) for 1h. The sections were counterstained in 4’,6-Diamidino-2-Phenylindole, Dihydrochloride (DAPI) (1:5000, Thermo Fisher Scientific, MA, USA) for 5 minutes and coverslips were mounted in ProLong™ Gold Antifade Mounting media (Thermo Fish Scientific, MA, USA). Images were captured using Leica SP8 Acousto-Optical Beam Splitter confocal laser scanning microscope attached to a Leica DM I8 inverted epifluorescence microscope.

#### Immunohistochemistry for neutrophil quantification in mouse kidney sections

Paraffin-embedded mouse kidney sections (5 μm) were dewaxed in xylene followed by rehydration in graded ethanol (100%, 90% and 70%). Following antigen retrieval in 10mM sodium citrate tribasic buffer(pH=6), endogenous peroxidase was blocked with 3% hydrogen peroxide (Merck, Darmstadt, Germany). The sections were blocked with 5% normal goat serum (Abcam, Cambridge, UK) followed by incubation with Neutrophil elastase antibody (1:200, Abcam, Cambridge, UK) overnight at 4 °C. After 3 washes, sections were incubated with HRP reagent (SignalStain® Boost Detection Reagent, Cell Signaling Technology, Leiden, Netherlands) for 30 minutes at room temperature. The positive staining was apparent with SignalStain® DAB Substrate Kit (Cell Signaling Technology, Leiden, Netherlands). Sections were counterstained with hematoxylin for 10 seconds and dehydrated with ethanol and xylene. The coverslips were mounted in ProLong™ Gold Antifade mounting media (Thermo Fish Scientific, MA, USA). The entire section was imaged using a Slide Scanner. Neutrophils were identified by their positive nuclear staining. The number of neutrophils was counted for each glomerulus and the mean glomerular neutrophil count for each mouse was analysed.

#### Mouse SDC4 and MMP2 and 9 ELISA

The concentration of SDC4 (ectodomain) in mouse plasma was quantified using a sandwich enzyme immunoassay (mouse SDC4 ELISA kit, CSB-EL020891MO, Cusabio, Houston, USA) according to the manufacturer’s instructions. Total MMP-2 Quantikine ELISA Kit, and total MMP-9 Immunoassay (R&D Systems, Minneapolis, USA) provide quantitation of total mouse MMP2 and 9 in serum and tissue homogenates. The assays were carried out according to the manufacturer’s instructions. The concentration of total MMP was normalized to total protein concentration.

#### Protein concentration

Total protein concentration was determined by Bicinchoninic Acid (BCA) assay (Thermo Fisher Scientific, MA, USA), carried out according to the manufacturer’s instructions.

#### RNA extraction

RNA was extracted from isolated glomeruli and a 2 mm^3^ chunk of kidney cortex using an RNeasy Mini kit (Qiagen, Manchester, UK) according to manufacturer’s instructions. Then, up to 2μg of total RNA was converted to cDNA by a high-capacity RNA to cDNA conversion kit (Applied Biosystems, Foster City, CA, USA), according to the manufacturer’s instructions.

#### Real-time PCR

The mRNA expression of relevant genes was quantified according to the manufacturer’s instructions for real-time PCR (StepOnePlus Real-Time PCR System; Applied Biosystems) using TaqMan primer probes (Life Technologies, ThermoFisher Scientific) detailed in supplementary Table 1. The 2^−ΔΔ^*^CT^* method was used to calculate the fold change, normalized to GAPDH.

#### Human SDC1, 4 and ANG2 ELISA

The levels of human SDC1, 4 (Human Syndecan-1, −4 DuoSet ELISA, R&D Systems, Minneapolis, USA), and ANGPT2 (Human Angiopoietin-2 ELISA Kit – Quantikine, R&D Systems, Minneapolis, USA) were determined in sepsis-AKI and control serum, according to the manufacturer’s instructions.

#### Human MMP2 and 9 activity assays

The activity of MMP2 (SensoLyte520 MMP-2 Plus Assay, AnaSpec Inc. & Eurogentec US, Fremont, USA) and 9 (Human Active MMP-9 Fluorokine E Kit, R&D Systems, Minneapolis, USA) was determined in sepsis-AKI and control serum, according to the manufacturer’s instructions.

## Statistical analysis

Data are expressed as the mean ± SEM. Normality was assessed using GraphPad Prism 5 Kolmogorov–Smirnov test. Normally distributed data was compared using *t*-tests for 2 groups and analysis of variance (ANOVA) for multiple groups. If one-way ANOVA indicated a significant difference, the Bonferroni post hoc test was used to assess differences between groups. Where normality could not be demonstrated, the Mann-Whitney test was used for comparing between 2 groups and Kruskal-Wallis was used for multiple groups. A p-value of <0.05 was considered to indicate statistical significance.

## Results

### Glomerular, peritubular and systemic eGlx are damaged in sepsis-AKI

To evaluate Glx damage in glomerular and peritubular capillaries and systemic circulation in sepsis-AKI, mice were injected with LPS and culled at 18 h post-LPS injection. Kidney sections, featuring glomerular and peritubular capillaries (Fig 1Ai, Bi), were labelled with lectin LEL and R18, identifying eGlx (green) and cell membrane (red), respectively. The anatomical distance between the 2 fluorescence peak profiles is a measure of eGlx depth (Fig 1Aii), as previously quantified.^22, 44^ Glomerular eGlx depth decreased by 0.48-fold in sepsis-AKI compared to control (Fig. 1Ai, Aiii, Aiv). Similarly, peritubular eGlx depth was reduced by 0.56-fold in sepsis-AKI compared to control (Fig. 1Bi, Bii, Biii). The reduction in renal eGlx depth was associated with a 4.56-fold increase in plasma SDC4 ectodomain, indicating enhanced systemic SDC4 shedding in sepsis-AKI (Fig. 1C).

### Elevated mRNA expression of eGlx components and endothelial inflammatory markers in the kidney is associated with increased sepsis-AKI markers

To determine changes in eGlx components and activated endothelial inflammation markers, TaqMan real-time PCR was performed on isolated glomeruli and whole kidney. The mRNA expression of SDC1 and 4 increased by 12.28- and 9.08-fold, respectively, in isolated glomeruli (Fig. 2A) and by 14.04- and 11.83-fold in whole kidney (Fig. 2B) in sepsis-AKI, suggesting possible compensatory feedback due to renal eGlx damage. ICAM and VCAM gene expression was upregulated by 12.47- and 4.56-fold in isolated glomeruli (Fig. 2A) and by 31.65- and 7.30-fold in whole kidney (Fig. 2B), indicating renal endothelial inflammation in sepsis-AKI. Additionally, these changes were associated with a 2.44- and 2.90-fold increase in serum creatinine and urea respectively (Fig. 2C, D).

**Figure 2.**
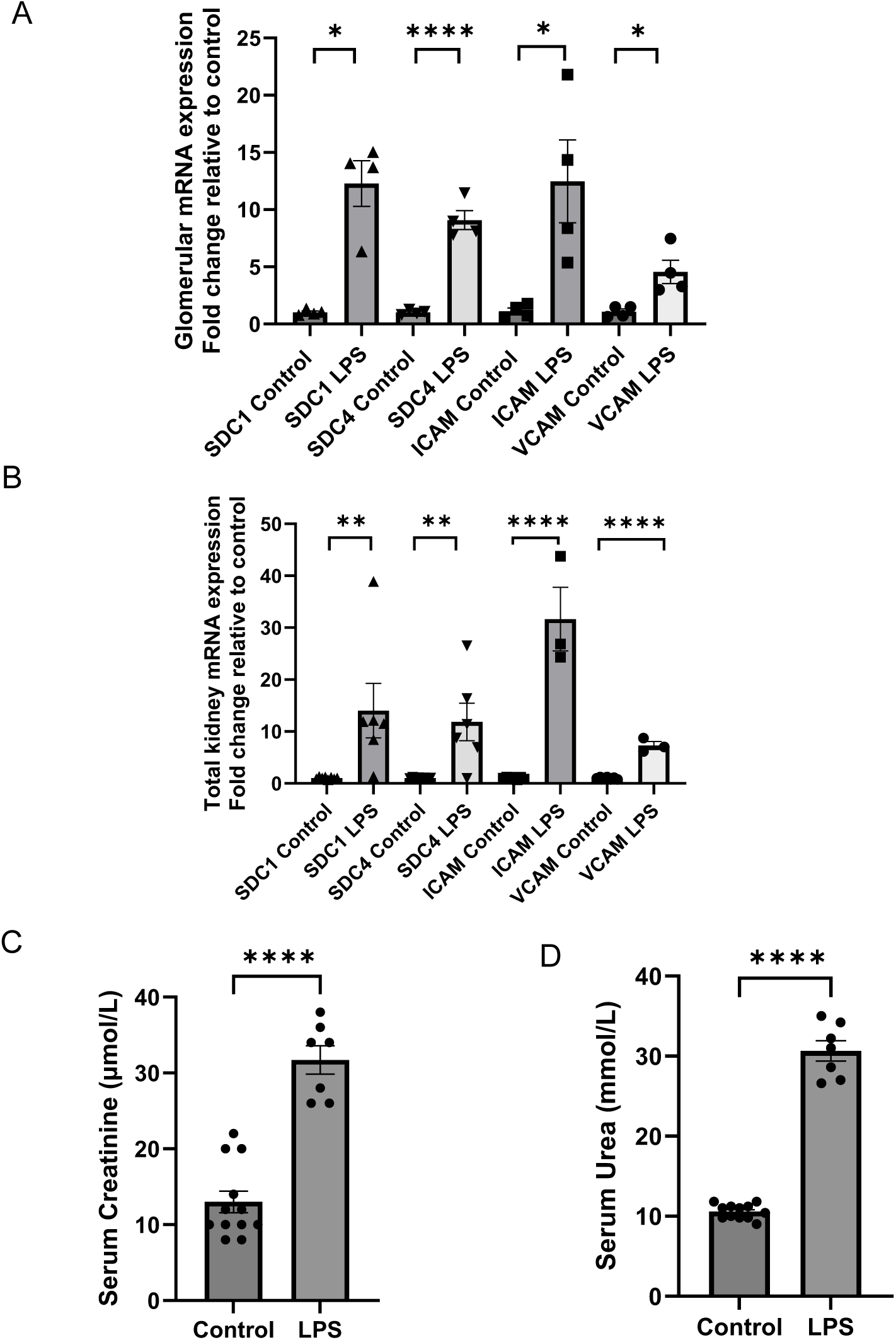
Elevated mRNA expression of eGlx components and endothelial inflammatory markers in the kidney is associated with increased sepsis-AKI markers. Isolated glomerular and total kidney mRNA expression of glycocalyx components SDC1 and SDC4 and endothelial adhesion markers ICAM-1 and VCAM-1 were determined. The 2^−ΔΔCT^ method of quantification was used to calculate fold change, normalised to GAPDH. (A) Isolated glomerular: SDC1 Control1.020 ± 0.1209 n=4 mice, 18h 12.28 ± 1.999, n=4 mice, *p=0.0286. SDC4: Control1.015 ± 0.1020 n=4 mice, 18h 9.082 ± 0.8257, n=4 mice, ****p=<0.0001. ICAM Control1.119 ± 0.2902 n=4 mice, 18h 12.47 ± 3.629, n=4 mice, *p=0.0206. VCAM Control1.083 ± 0.2388 n=4 mice, 18h 4.555 ± 1.024, n=4 mice, *p=0.0164. (B) Total kidney SDC1 Control1.021±0.07570 n=8 mice, 18h 14.04 ± 5.248 n=6 mice, **p=0.0013. SDC4: Control1.006 ± 0.04003 n=8 mice, 18h 11.83 ± 3.594 n=6 mice **p=0.0042. ICAM: Control1.020 ± 0.07864 n=7 mice, 18h 31.65 ± 6.114 n=3 mice ****p<0.0001. VCAM: Control 1.016 ± 0.07115 n=7 mice, 18h 7.295 ± 0.7545 n=3 mice ****p<0.0001. (C) Serum creatinine control 13.00 ± 1.425 n=12 mice, 18h LPS 31.71 ± 1.874 n=7 mice, ****p<0.0001. (B) Serum urea control 10.57 ± 0.2580 n=12 mice, 18h LPS 30.66 ± 1.272 n=7 mice, ***p<0.0001. Each dot on the graph represents a mouse. Data are expressed as the mean ± SEM. An unpaired t-test was used if the data were normally distributed. If normality was not achieved, Mann Whitney test was used for statistical analysis.

### Renal eGlx damage is associated with increased sheddase MMP9 and reduced MMP2 levels in sepsis-AKI

To determine if damage to the renal eGlx was associated with an increase in shedding enzymes (also known as sheddases), the expression levels of MMP2 and MMP9 were examined. Immunofluorescence staining revealed that MMP9 is mainly expressed in the glomeruli (identified by nephrin labelling), with very low expression in control mice but increased expression in LPS-treated mice (Fig. 3Ai). Furthermore, co-staining of MMP9 with markers of glomerular cells: endomucin for endothelial cells, nephrin for podocytes and PDGFR-β for mesangial cells, and calgranulin A for neutrophils, showed evidence of colocalisation with neutrophils, endothelial cells and mesangial cells but not with podocytes (Fig. 3Aii-iii). MMP9 protein expression was increased in isolated glomeruli and the total kidney as analysed by western blot in sepsis-AKI (Fig. 3B). MMP9 mRNA expression increased by 9.50- and 3.64-fold in isolated glomeruli and the total kidney, respectively in sepsis-AKI (Fig. 3C). Additionally, serum MMP9 levels were elevated by 4.45-fold in sepsis-AKI (Fig. 3D). In contrast, MMP2 mRNA expression decreased by 0.44- and 0.21-fold in isolated glomeruli and the total kidney, respectively, in LPS compared to controls (Fig. 3E). There was no significant change in serum MMP2 levels in LPS compared to the control group (Fig. 3F).

**Figure 3.**
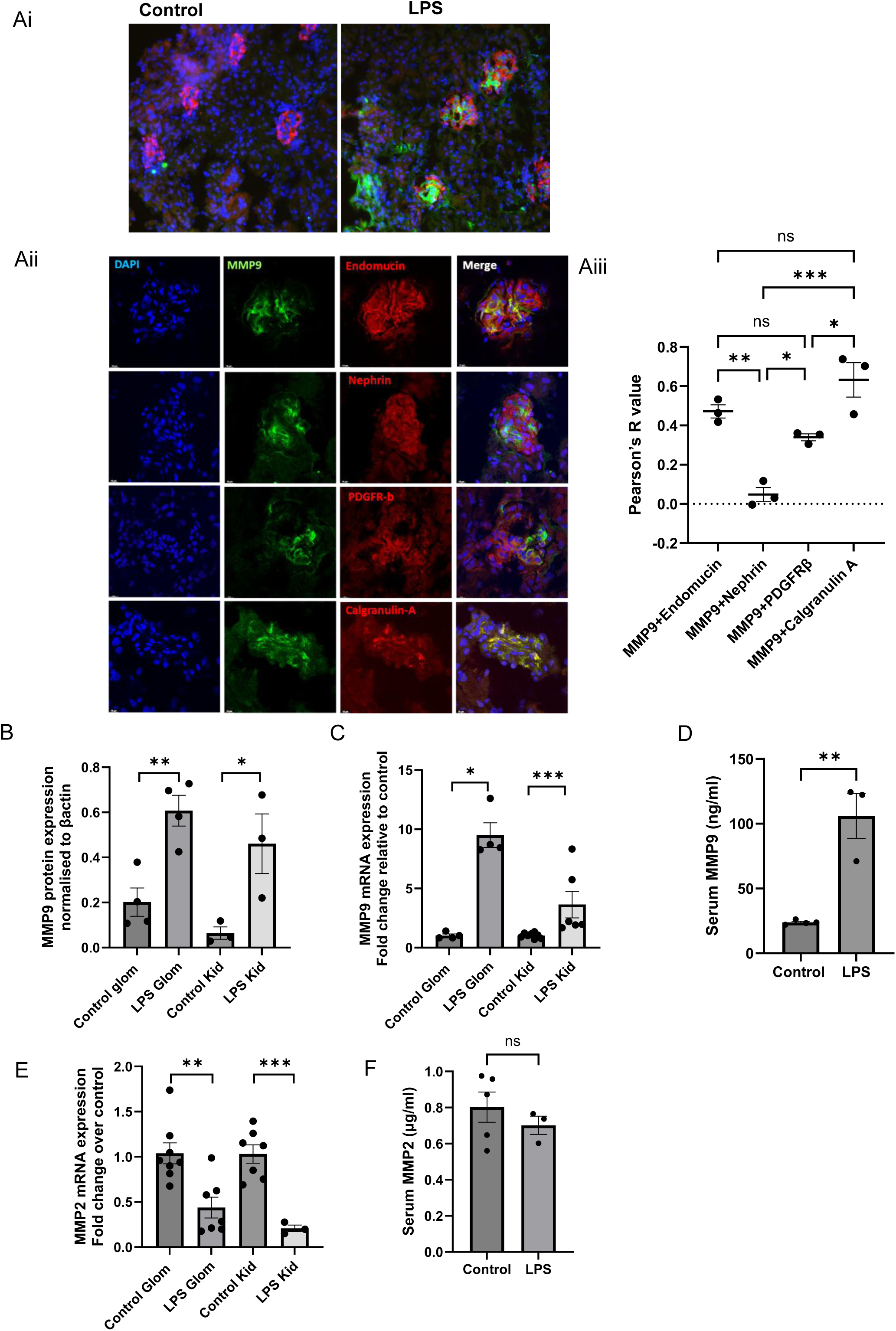
Renal eGlx damage is associated with increased sheddase MMP9 and reduced MMP2 levels in sepsis-AKI. (Ai) Immunofluorescence staining of MMP9 (green) and nephrin (red) on kidney sections from control and LPS injected mice. (Aii) Colocalisation of MMP9 expression (green) with endothelial marker, endomucin (red), podocyte marker, nephrin (red), mesangial marker, platelet-derived growth factor (PDGFβ) (red), neutrophil marker, calgranulin A (red) and nuclei marker DAPI (blue). (Aiii) Pearson R correlation quantifies the degree of correlation between MMP9 expression and the markers. Western blot densitometry compares MMP9 to β-actin expression in (B) isolated glomeruli: control 0.2018 ± 0.06282 n=4, LPS 0.6074 ± 0.06830, n=4, **p=. 0.0047 and total kidney: control 0.06435 ± 0.02739 n=3, LPS 0.4608 ± 0.1324 n=3, *p=0.0427. The 2^−ΔΔCT^ method of quantification was used to calculate the fold change in mRNA expression, normalised to GAPDH (C) Isolated glomerular MMP9: Control1.025 ± 0.1366 n=4 mice, LPS 9.503 ± 1.036, n=4 mice, *p=0.0286; total kidney MMP9: Control1.025 ± 0.07955 n=8 mice, 18h 3.642 ± 1.126 n=3 mice, ***p=0.0007. (D) MMP9 level in serum in control 23.84 ± 0.8767 n= 4 mice, LPS 106.0 ± 17.45 n=3 mice, **p=0.0025. (E) Isolated glomerular MMP2: Control1.038 ± 0.1153 n=8 mice, LPS 0.4372 ± 0.1151, n=7 mice, **p=0.0028; total kidney MMP2: Control1.031 ± 0.1017 n=7 mice, LPS 0.2075 ± 0.03682 n=3 mice, ***p=0.0010. (F) MMP2 level in serum control 0.8026 ± 0.08399 n=5 mice, LPS 0.7014 ± 0.05039 n=3 mice, ns non-significant. Each dot on the graph represents a mouse. Data are expressed as the mean ± SEM. One-way ANOVA with Bonferroni’s multiple comparison test or unpaired t-test, as appropriate, if data were not normally distributed, Kruskal-Wallis test multiple comparison or Mann Mann-Whitney test, as appropriate, was used for statistical analysis.

### Blockade of MMP2 and 9 restores glomerular, peritubular and systemic eGlx in sepsis-AKI

To evaluate whether inhibition of MMP2 and 9 protects renal and systemic eGlx, mice were injected with 5mg/kg MMPI 1h before LPS injection (Fig4A). We focused our analysis on MMP9, given the increase in MMP9 and not MMP2 observed above. Peak-to-peak analysis was carried out to assess glomerular and peritubular eGlx depth, as above. LPS caused a 0.86- and 0.74-fold reduction in glomerular and peritubular eGlx depth, respectively, which were restored to control levels by MMPI (Fig. 4A, B). Similarly, LPS induced 6.20-fold increase in serum SDC4, which was attenuated by 0.78-fold with MMPI treatment (Fig. 4C). We confirmed that LPS enhanced glomerular, kidney and circulating MMP9 expression by 13.96-, 5.05- and 5.26-fold (Fig. 4D-F). MMPI reduced glomerular MMP9 levels by 0.25-fold when compared to LPS-treated mice (Fig. 4D), but didn’t significantly lower MMP9 in serum and kidney (Fig. 4E, F). Glomerular MMP9 negatively correlated with glomerular eGlx in sepsis-AKI (Fig. 4G). There was a significant reduction in blood pressure in sepsis-AKI. MMPI had no significant effect on blood pressure measurement in sepsis-AKI (Sup. Fig.1A, B). There was also no significant change in body weight in sepsis-AKI mice when compared to control. Equally, MMPI did not affect body weight in sepsis-AKI over the 18h period (Sup. Fig.1C).

**Figure 4.**
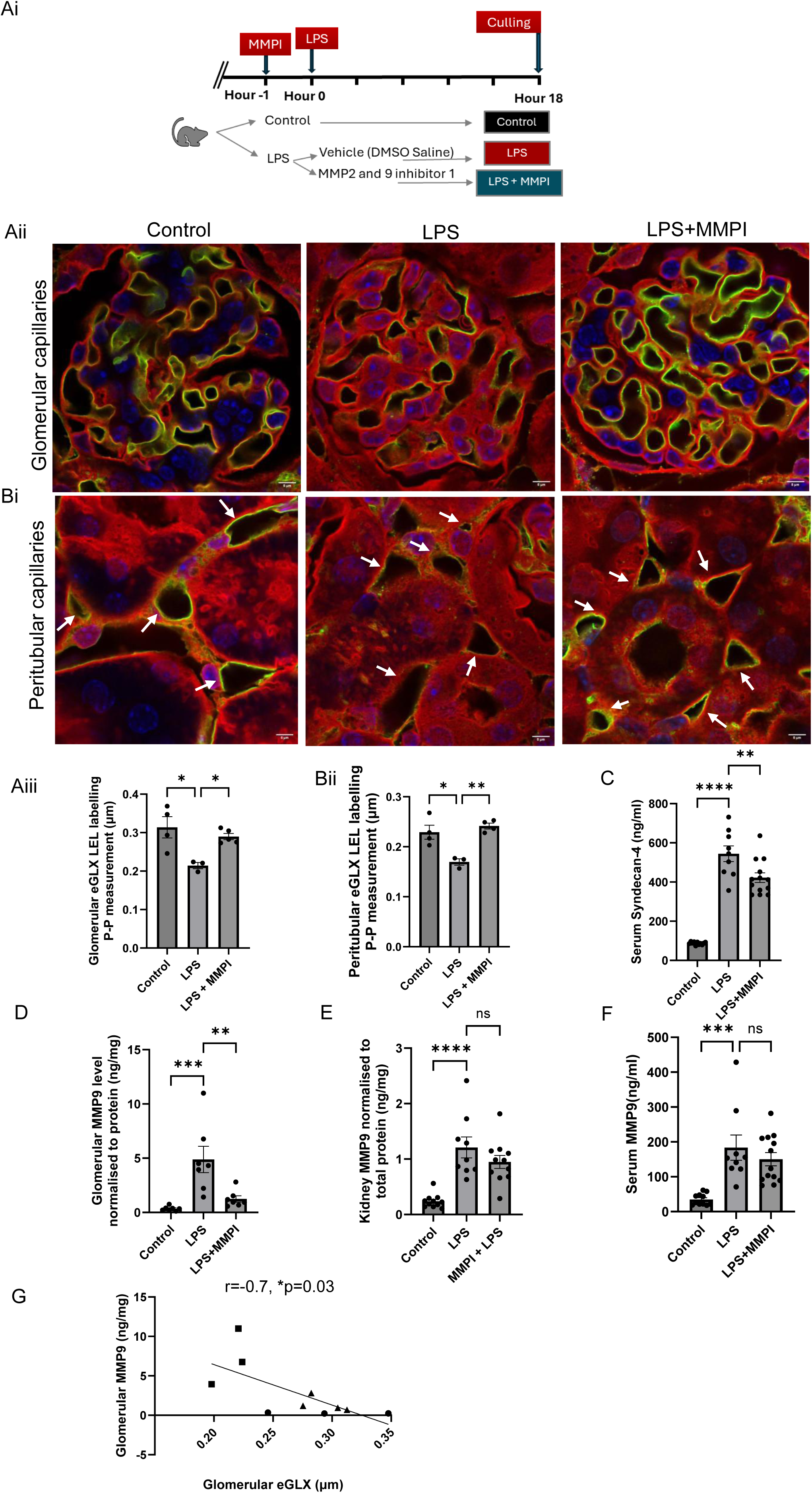
Blockade of MMP 2 and 9 restores glomerular, peritubular and systemic eGlx in sepsis-AKI. (Ai) Schematic overview of the timeline: mice were injected with 5mg/kg MMPI or vehicle 1h prior to 10mg/kg LPS injection. Control mice received saline as vehicle. All mice were culled 18h post LPS injection. Kidney sections were stained with Lycopersicon esculentum (Tomato) Lectin (LEL) and endothelial membrane label R18. Representative image shows (Aii) glomerular and (Bi) peritubular capillaries labelled red (R18) and the luminal endothelial glycocalyx labelled green (LEL). Peak to Peak assessment of the eGlx depth post-LPS injections (Aiii) glomerular capillaries control 0.3139 ± 0.02771 n=4 mice, LPS 0.2142 ± 0.008135 n=3 mice, MMPI+LPS 0.2897± 0.008016, n=5 mice, *p<0.05. (Bii) Peak to Peak assessment of the peritubular eGlx depth, control 0.2291± 0.01393 n=4 mice, LPS 0.1695±0.006473 n=3 mice, MMPI+LPS 0.2416.± 0.005814 n=4 mice *p<0.05, **p<0.005. (C) Serum SDC4, control 87.83± 2.268 n=11 mice, LPS 544.3±39.81 n=9 mice, MMPI+LPS 422.3.± 24.88 n=13 mice **p=0.0043, ****<0.0001. MMP9 expression in (D) isolated glomeruli, control 0.3498 ± 0.07711 n= 7mice, LPS 4.882 ± 1.209 n=7mice, MMPI+LPS 1.238 ± 0.2883 n=7mice **p<0.005, ***p=0.0006 (E) kidney, control 0.2394 ± 0.03747 n= 11mice, LPS 1.209 ± 0.1891 n=9mice, MMPI+LPS 0.9482 ± 0.1171 n=11mice ****p<0.0001, ns - non significant (F) serum, control 34.89 ± 4.980 n= 11mice, LPS 183.6 ± 36.30 n=9mice, MMPI+LPS 150.3 ± 18.91 n=13mice ***p=0.0001, ns - non significant (G) Correlation between glomerular MMP9 concentration and glomerular eGlx, circle is control, square is LPS and triangle is MMPI+LPS, Pearson r= −0.68, *p=<0.05. Each dot on the graph represents a mouse. Data are expressed as the mean ± SEM. One-way ANOVA with Bonferroni’s multiple comparison test was used for statistical analysis.

### MMP2 and 9 inhibition attenuates neutrophil count and ICAM expression in the kidney in sepsis-AKI

To determine whether sepsis-AKI is associated with neutrophil trafficking and whether MMP2 and 9 blockade reduces influx, immunohistochemistry was performed on kidney sections using a neutrophil elastase antibody. Sepsis-AKI was associated with an 11.89-fold increase in average neutrophil count in the glomeruli, which was attenuated by 0.67-fold in the MMPI-treated group (Fig. 5Ai, Aii). In addition, renal ICAM-1 expression was increased by 22.77-fold in sepsis-AKI, and attenuated by 0.53-fold in the presence of the MMPI (Fig. 5B), confirming reduced leucocyte trafficking and inflammation in the kidney.

**Figure 5.**
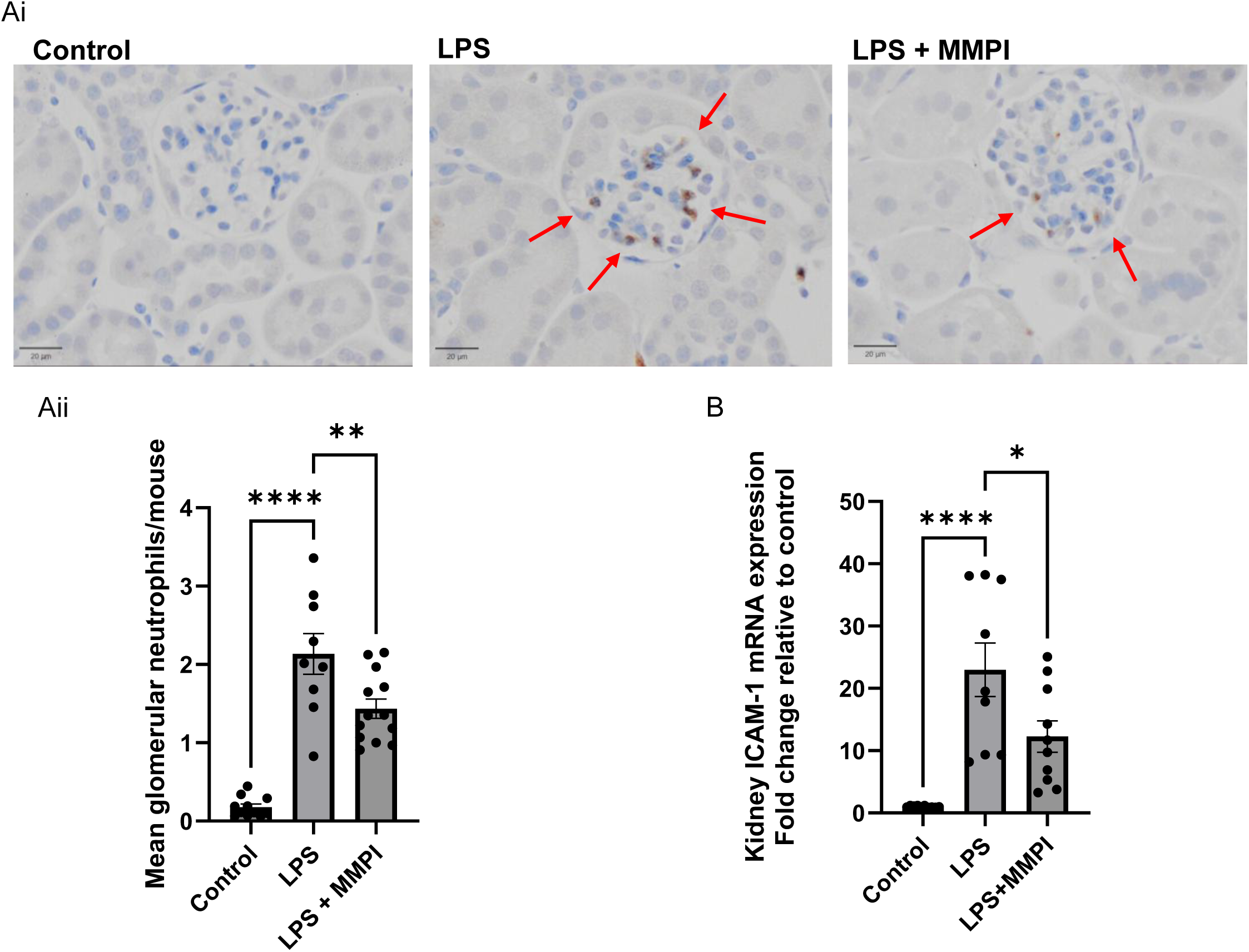
MMP2 and 9 inhibition attenuates neutrophil count and ICAM expression in the kidney in sepsis-AKI. A. Neutrophil elastase-positive stained neutrophils, indicated by red arrows on kidney sections: control, LPS and LPS+MMPI treated mice. Aii. The average neutrophil count for all glomeruli in each mouse is control 0.1796±0.03857,n=11mice, LPS 2.136±0.2601, n=9 mice, LPS+MMPI 1.435±0.1224, n-13 mice, ****p<0.0001,**p=0.0055. B. Kidney ICAM-1 gene expression relative to GAPDH for each mouse is control 1.009±0.04637, n=10 mice, LPS 22.97±4.303, n=9 mice, LPS+MMPI 12.26±2.522, n=10 mice, ****p<0.0001, *p=0.0228. Data are expressed as the mean ± SEM. One-way ANOVA with Bonferroni’s or Kruskal-Wallis multiple comparison test, as appropriate, was used for statistical analysis.

### Increased MMP9 activity, eGlx damage and endothelial activation in sepsis-AKI in humans

To determine the relevance of the sheddases (MMP2 and 9) in eGlx damage in human sepsis-AKI, an observational cohort study was performed on 34 participants: 16 sepsis-AKI patients and 18 age-matched healthy controls (Table 1). Sepsis-AKI patients had a 2.79-,1.5- and 2.11-fold increase in serum MMP9 activity, SDC1 shedding and ANGPT2 concentration, respectively, but no change in MMP2 activity or SDC4 shedding (Sup Fig. 2A-B). To assess whether sepsis-AKI serum has a direct and detrimental effect on renal endothelium, human GEnC Glx was prelabelled with ManNAz and exposed to 10% sepsis-AKI or control serum. Exposure to sepsis-AKI serum resulted in a 0.68-fold loss of ManNAz-labelled eGlx when compared to human GEnC treated with control serum, demonstrating a direct and detrimental effect of human sepsis-AKI serum on human renal eGlx (Fig. 6Ei-iii).

**Figure 6.**
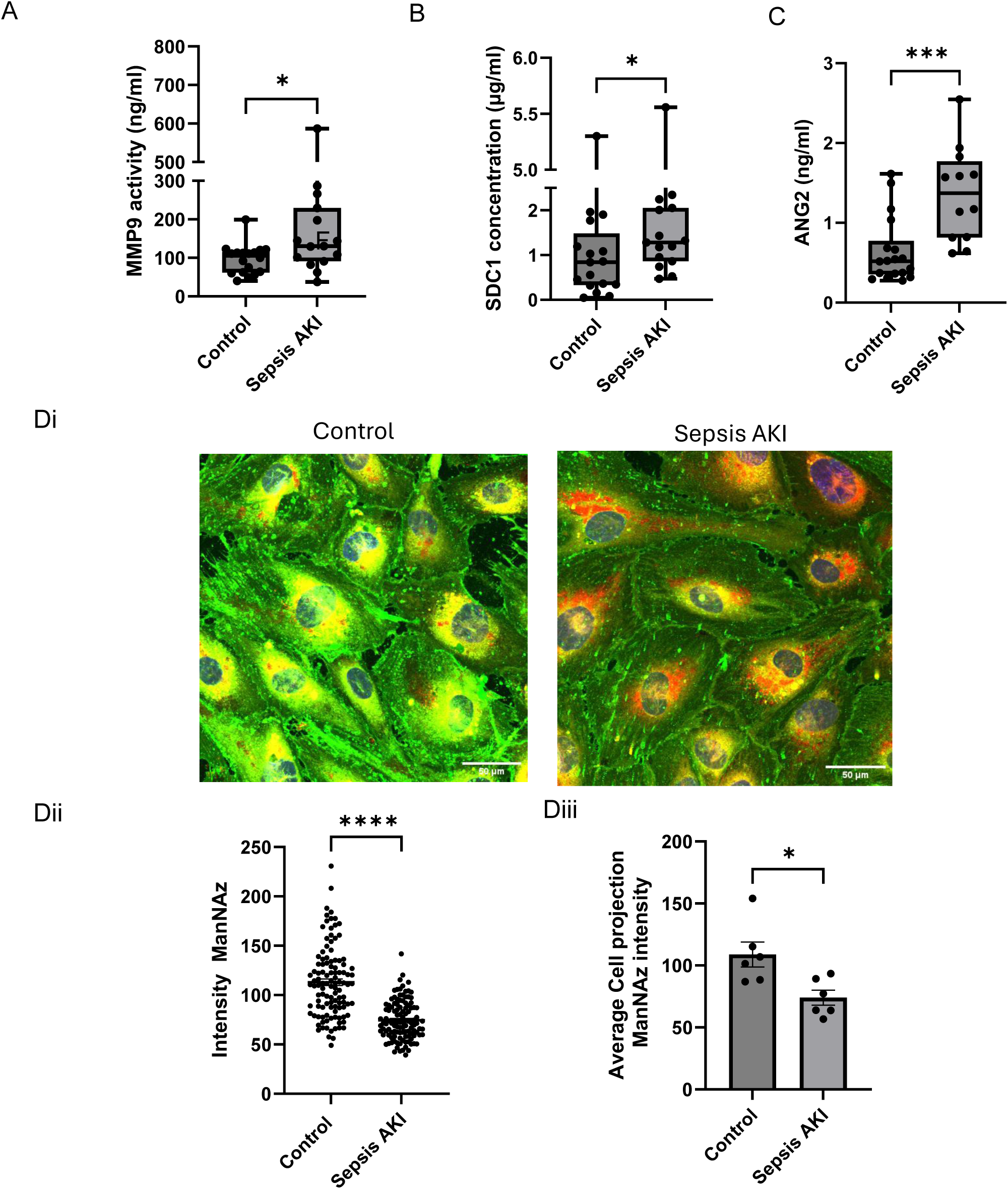
Increased MMP9 activity, eGlx damage and endothelial activation in sepsis-AKI in humans. (A) MMP9 activity: control 0.9588 ± 0.08998 n=18, sepsis-AKI:2.674 ± 0.9966 n=15 *p=0.0184, (B) SDC1 shedding: control 1.074 ± 0.3036 n=17, sepsis-AKI 1.611 ± 0.3214 n=15 *p=0.0244, (C) ANG2 expression control 0.6414 ± 0.09745 n=18, sepsis-AKI 1.357 ± 0.1708 n=12 ***p= 0.0004. (Di) Click-iT ManNAz-‘labelled’ human GEnC were exposed to 10% human serum from control and sepsis patients for 2h. (Dii) The ManNAz fluorescence intensity was quantified control 113.0 ± 3.391, sAKI 74.45 ± 1.821, ****p<0.0001. (Diii) The average cell projection ManNAz intensity control 108.8 ± 10.08, sAKI 74.03 ± 6.121, n=6, *p=0.0145. Each dot on the graph represents a participant. Data are expressed as box plots showing median [IQR]. Some data are expressed as mean ± SEM. Unpaired t-test was used for statistical analysis. Student t test or Mann-Whitney test was used, dependent on whether the data was normally distributed.

## Discussion

This study demonstrates that renal and systemic eGlx are impaired in sepsis-AKI. This impairment is likely to be mediated by sheddase MMP9, which results in eGlx shedding and contributes to microvascular dysfunction. We also confirmed the significance of MMP2 and/or 9 activity and microvascular dysfunction in sepsis-AKI in humans, indicating that MMP2 and 9 blockade may be a potential therapeutic target for protecting eGlx and restoring renal microvascular damage in sepsis-AKI.

To our knowledge, this is the first study demonstrating eGlx damage in renal microvascular beds, specifically in glomerular and peritubular capillaries in sepsis-AKI. We also showed increased circulating SDC4 in mice, indicating systemic eGlx damage in sepsis-AKI. Our findings, which demonstrate renal eGlx damage in mouse sepsis-AKI, align with previous studies showing glomerular glycocalyx loss (stained with WGA) in sepsis-AKI at 24 hours post-LPS injection^7^ and early changes in the glomerular filtration barrier-associated glycocalyx in caecal ligation and puncture-induced kidney damage.^46, 47^ However, none of the studies specifically demonstrated *endothelial* Glx, as WGA staining non-selectively binds to carbohydrate residues present on multiple glomerular cell types in the kidney.

Previously, we identified MMP2 and 9-mediated eGlx shedding as a key mechanism in GEnC Glx damage and microvascular dysfunction in diabetes and a RAAS activation model of kidney injury. In this study, we sought to investigate whether this mechanism contributes to renal and systemic microvascular endothelial cell injury in a mouse model of sepsis-AKI. We established that MMP9 but not MMP2 levels were increased in sepsis-AKI. MMP9 expression was elevated in both isolated glomeruli and kidney at protein and mRNA levels, although its expression was predominantly in the glomeruli. In the glomeruli, MMP9 expression colocalises strongly with calgranulin A, a neutrophil marker, endomucin, an endothelial marker and PDGFβ, mesangial cell marker, but not with nephrin, podocyte marker. This demonstrates that neutrophils, GEnC and mesangial cells are likely sources of MMP9 in glomeruli. Our data are consistent with other AKI models, which demonstrate an increase in MMP9 in the kidney in cisplatin-induced AKI^48^ and ischemia-reperfusion injury-induced AKI.^49^ In addition, Caron A *et al*., demonstrated increased MMP9 activity in isolated renal endothelial cell fractions in ischemia-AKI,^50^ substantiating the GEnC expression of MMP9 in our model.

We then sought to determine whether blocking MMP2 and 9 would restore eGlx and protect from renal microvasculature dysfunction. Inhibition of MMP2 and 9 with a potent gelatinase inhibitor, used previously,^22, 35^ restored eGlx on glomerular and peritubular capillaries to control levels and reduced systemic SDC4 shedding. We also confirmed a significant reduction in MMP9 levels in glomeruli. This was accompanied by reduced leucocyte trafficking in the glomeruli in sepsis-AKI. In line with this, studies have found that inhibition of MMP2 and 9 with minocycline diminished leucocyte accumulation and renal microvascular permeability following ischemia.^31^ Moreover, minocycline had anti-inflammatory and anti-apoptotic effects, reduced ICAM-1 expression and serum creatinine in a rat model of ischemic renal injury.^51^ Similarly, inhibition of MMP2 and 9 with tetracycline doxycycline had anti-inflammatory effects, reduced leucocyte infiltration^52^ and reduced kidney damage in cisplatin-induced AKI.^48^ MMP9 deletion plays a key role in reducing neutrophil infiltration and protecting from other organ injury e.g., in ischemia-reperfusion-induced myocardial injury^53^ and liver ischemia-reperfusion injury.^33^ Future work will look into using MMP9-deficient mice to confirm whether MMP9 deficiency protects against kidney injury in sepsis-AKI.

To determine the relevance of MMP9 activity, eGlx shedding and endothelial damage in sepsis-AKI in humans, an observational cohort study and a sepsis-AKI in vitro study were performed. We observed an increase in MMP9 activity in sepsis-AKI patients. This was accompanied by an increase in SDC1 shedding and enhanced endothelial damage marker ANGPT2. The latter is an antagonist of endothelial-stabilising receptor Tie2, secreted by endothelial cells and it promotes vascular permeability and neutrophil recruitment and infiltration into the endothelium.^54^ Kumpers group demonstrated that ANGPT2 mediates eGlx damage, vascular leakage and leucocyte extravasation through heparanase,^54, 55^ suggesting that heparanase might also be playing a key role in eGlx damage in sepsis-AKI. In addition, we confirmed the direct and detrimental effect of (MMP9)/sepsis-AKI serum on renal microvasculature in humans. This demonstrates that blockade of MMP 2 and 9 might be a potential therapeutic target for protecting eGlx and restoring renal microcirculation in sepsis-AKI.

We acknowledge that there are some limitations in this study. We have focused on MMPI as a preventative measure and it would be informative to know whether MMPI, given therapeutically, after initiation of sepsis, would restore eGlx and protect from microvascular damage in sepsis-AKI. Ideally, a time course identifying the optimum time to block MMP9 in sepsis-AKI is warranted. All mice in this study are male, because studies have shown that sex hormones have an impact on immune cells, leading to dimorphic response to LPS.^56^ To avoid sexual dimorphism being a confounder, only male mice were included in this study. A potent MMP2 and 9 inhibitor is used in this study, although effective at blocking MMP9 in the glomeruli, it was not effective at reducing MMP9 levels in the serum and kidney tissue. Of note, in the in vivo study, total MMP9 levels were quantified rather than MMP9 activity, which itself might not capture the full extent of the MMP9 inhibition.

In summary, our data reveal a crucial role of MMP2 and 9 in renal and systemic eGlx damage in sepsis-AKI and that blockade of MMP2 and 9 restores renal and systemic eGlx and protects from neutrophil trafficking in the glomeruli. The results advance our understanding of the mechanism of renal eGlx damage and renal microvascular dysfunction in sepsis-AKI.

## Acknowledgements

We would like to thank Dr Joe Roe, the Named Veterinary Surgeon at the University of Bristol, for his help and support with the sepsis-AKI animal model and associated animal welfare. We would also like to thank Dr Amulic for his kind gift of the Calgranulin A antibody.

## References

1. Peerapornratana S, Manrique-Caballero CL, Gomez H, et al. Acute kidney injury from sepsis: current concepts, epidemiology, pathophysiology, prevention and treatment. Kidney Int 2019; 96: 1083–1099.

2. Peerapornratana S, Priyanka P, Wang S, et al. Sepsis-Associated Acute Kidney Disease. Kidney Int Rep 2020; 5: 839–850.

3. Zarbock A, Nadim MK, Pickkers P, et al. Sepsis-associated acute kidney injury: consensus report of the 28th Acute Disease Quality Initiative workgroup. Nat Rev Nephrol 2023; 19: 401–417.

4. Bellomo R, Kellum JA, Ronco C, et al. Acute kidney injury in sepsis. Intensive Care Med 2017; 43: 816–828.

5. Molema G, Zijlstra JG, van Meurs M, et al. Renal microvascular endothelial cell responses in sepsis-induced acute kidney injury. Nat Rev Nephrol 2022; 18: 95–112.

6. Molitoris BA. Therapeutic translation in acute kidney injury: the epithelial/endothelial axis. J Clin Invest 2014; 124: 2355–2363.

7. Xu C, Chang A, Hack BK, et al. TNF-mediated damage to glomerular endothelium is an important determinant of acute kidney injury in sepsis. Kidney Int 2014; 85: 72–81.

8. Ramnath R, Foster RR, Qiu Y, et al. Matrix metalloproteinase 9-mediated shedding of syndecan 4 in response to tumor necrosis factor alpha: a contributor to endothelial cell glycocalyx dysfunction. FASEB J 2014; 28: 4686–4699.

9. Ermert K, Buhl EM, Klinkhammer BM, et al. Reduction of Endothelial Glycocalyx on Peritubular Capillaries in Chronic Kidney Disease. Am J Pathol 2023; 193: 138–147.

10. Nelson A, Berkestedt I, Schmidtchen A, et al. Increased levels of glycosaminoglycans during septic shock: relation to mortality and the antibacterial actions of plasma. Shock 2008; 30: 623–627.

11. Banerjee S, Mohammed A, Wong HR, et al. Machine Learning Identifies Complicated Sepsis Course and Subsequent Mortality Based on 20 Genes in Peripheral Blood Immune Cells at 24 H Post-ICU Admission. Front Immunol 2021; 12: 592303.

12. Cox LA, van Eijk LT, Ramakers BP, et al. Inflammation-induced increases in plasma endocan levels are associated with endothelial dysfunction in humans in vivo. Shock 2015; 43: 322–326.

13. Scherpereel A, Depontieu F, Grigoriu B, et al. Endocan, a new endothelial marker in human sepsis. Crit Care Med 2006; 34: 532–537.

14. Chelazzi C, Villa G, Mancinelli P, et al. Glycocalyx and sepsis-induced alterations in vascular permeability. Crit Care 2015; 19: 26.

15. Schmidt EP, Overdier KH, Sun X, et al. Urinary Glycosaminoglycans Predict Outcomes in Septic Shock and Acute Respiratory Distress Syndrome. Am J Respir Crit Care Med 2016; 194: 439–449.

16. Inkinen N, Pettila V, Lakkisto P, et al. Association of endothelial and glycocalyx injury biomarkers with fluid administration, development of acute kidney injury, and 90-day mortality: data from the FINNAKI observational study. Ann Intensive Care 2019; 9: 103.

17. Singh A, Satchell SC, Neal CR, et al. Glomerular endothelial glycocalyx constitutes a barrier to protein permeability. J Am Soc Nephrol 2007; 18: 2885–2893.

18. Singh A, Friden V, Dasgupta I, et al. High glucose causes dysfunction of the human glomerular endothelial glycocalyx. Am J Physiol Renal Physiol 2011; 300: F40–48.

19. Singh A, Ramnath RD, Foster RR, et al. Reactive oxygen species modulate the barrier function of the human glomerular endothelial glycocalyx. PLoS One 2013; 8: e55852.

20. Desideri S, Onions KL, Qiu Y, et al. A novel assay provides sensitive measurement of physiologically relevant changes in albumin permeability in isolated human and rodent glomeruli. Kidney Int 2018; 93: 1086–1097.

21. Crompton M, Ferguson JK, Ramnath RD, et al. Mineralocorticoid receptor antagonism in diabetes reduces albuminuria by preserving the glomerular endothelial glycocalyx. JCI Insight 2023; 8: e154164.

22. Ramnath RD, Butler MJ, Newman G, et al. Blocking matrix metalloproteinase-mediated syndecan-4 shedding restores the endothelial glycocalyx and glomerular filtration barrier function in early diabetic kidney disease. Kidney Int 2020; 97: 951–965.

23. Salmon AH, Ferguson JK, Burford JL, et al. Loss of the endothelial glycocalyx links albuminuria and vascular dysfunction. J Am Soc Nephrol 2012; 23: 1339–1350.

24. Gamez M, Ramnath RD, Butler MJ, et al. The glomerular endothelial glycocalyx as a therapeutic target in proteinuric kidney disease. Nat Rev Nephrol 2025.

25. Garsen M, Benner M, Dijkman HB, et al. Heparanase Is Essential for the Development of Acute Experimental Glomerulonephritis. Am J Pathol 2016; 186: 805–815.

26. Rops AL, Loeven MA, van Gemst JJ, et al. Modulation of heparan sulfate in the glomerular endothelial glycocalyx decreases leukocyte influx during experimental glomerulonephritis. Kidney Int 2014; 86: 932–942.

27. Devi S, Li A, Westhorpe CL, et al. Multiphoton imaging reveals a new leukocyte recruitment paradigm in the glomerulus. Nat Med 2013; 19: 107–112.

28. Babickova J, Klinkhammer BM, Buhl EM, et al. Regardless of etiology, progressive renal disease causes ultrastructural and functional alterations of peritubular capillaries. Kidney Int 2017; 91: 70–85.

29. Schmidt EP, Yang Y, Janssen WJ, et al. The pulmonary endothelial glycocalyx regulates neutrophil adhesion and lung injury during experimental sepsis. Nat Med 2012; 18: 1217–1223.

30. Mulivor AW, Lipowsky HH. Role of glycocalyx in leukocyte-endothelial cell adhesion. Am J Physiol Heart Circ Physiol 2002; 283: H1282–1291.

31. Sutton TA, Kelly KJ, Mang HE, et al. Minocycline reduces renal microvascular leakage in a rat model of ischemic renal injury. Am J Physiol Renal Physiol 2005; 288: F91–97.

32. Asahi M, Asahi K, Jung JC, et al. Role for matrix metalloproteinase 9 after focal cerebral ischemia: effects of gene knockout and enzyme inhibition with BB-94. J Cereb Blood Flow Metab 2000; 20: 1681–1689.

33. Hamada T, Fondevila C, Busuttil RW, et al. Metalloproteinase-9 deficiency protects against hepatic ischemia/reperfusion injury. Hepatology 2008; 47: 186–198.

34. Yazdan-Ashoori P, Liaw P, Toltl L, et al. Elevated plasma matrix metalloproteinases and their tissue inhibitors in patients with severe sepsis. J Crit Care 2011; 26: 556–565.

35. Butler MJ, Ramnath R, Kadoya H, et al. Aldosterone induces albuminuria via matrix metalloproteinase-dependent damage of the endothelial glycocalyx. Kidney Int 2019; 95: 94–107.

36. Li H, Mittal A, Makonchuk DY, et al. Matrix metalloproteinase-9 inhibition ameliorates pathogenesis and improves skeletal muscle regeneration in muscular dystrophy. Human molecular genetics 2009; 18: 2584–2598.

37. Yamaguchi M, Jadhav V, Obenaus A, et al. Matrix metalloproteinase inhibition attenuates brain edema in an in vivo model of surgically-induced brain injury. Neurosurgery 2007; 61: 1067–1075; discussion 1075-1066.

38. Tamura Y, Watanabe F, Nakatani T, et al. Highly selective and orally active inhibitors of type IV collagenase (MMP-9 and MMP-2): N-sulfonylamino acid derivatives. J Med Chem 1998; 41: 640–649.

39. Vandenbroucke RE, Libert C. Is there new hope for therapeutic matrix metalloproteinase inhibition? Nat Rev Drug Discov 2014; 13: 904–927.

40. Desideri S, Onions KL, Qiu Y, et al. A novel assay provides sensitive measurement of physiologically relevant changes in albumin permeability in isolated human and rodent glomeruli. Kidney international 2018.

41. Singer M, Deutschman CS, Seymour CW, et al. The Third International Consensus Definitions for Sepsis and Septic Shock (Sepsis-3). JAMA 2016; 315: 801–810.

42. Kellum JA, Lameire N, Group KAGW. Diagnosis, evaluation, and management of acute kidney injury: a KDIGO summary (Part 1). Crit Care 2013; 17: 204.

43. Goyard D, Diriwari PI, Berthet N. Metabolic labelling of cancer cells with glycodendrimers stimulate immune-mediated cytotoxicity. RSC Med Chem 2022; 13: 72–78.

44 Crompton M, Ferguson JK, Ramnath RD, et al. Mineralocorticoid receptor antagonism in diabetes reduces albuminuria by preserving the glomerular endothelial glycocalyx. JCI Insight 2023; 8.

45. Knackstedt SL, Georgiadou A, Apel F, et al. Neutrophil extracellular traps drive inflammatory pathogenesis in malaria. Sci Immunol 2019; 4.

46. Adembri C, Sgambati E, Vitali L, et al. Sepsis induces albuminuria and alterations in the glomerular filtration barrier: a morphofunctional study in the rat. Crit Care 2011; 15: R277.

47. Lygizos MI, Yang Y, Altmann CJ, et al. Heparanase mediates renal dysfunction during early sepsis in mice. Physiol Rep 2013; 1: e00153.

48. Nakagawa T, Kakizoe Y, Iwata Y, et al. Doxycycline attenuates cisplatin-induced acute kidney injury through pleiotropic effects. Am J Physiol Renal Physiol 2018; 315: F1347–F1357.

49. Kunugi S, Shimizu A, Kuwahara N, et al. Inhibition of matrix metalloproteinases reduces ischemia-reperfusion acute kidney injury. Lab Invest 2011; 91: 170–180.

50. Caron A, Desrosiers RR, Beliveau R. Ischemia injury alters endothelial cell properties of kidney cortex: stimulation of MMP-9. Exp Cell Res 2005; 310: 105–116.

51. Kelly KJ, Sutton TA, Weathered N, et al. Minocycline inhibits apoptosis and inflammation in a rat model of ischemic renal injury. Am J Physiol Renal Physiol 2004; 287: F760–766.

52. Mulivor AW, Lipowsky HH. Inhibition of glycan shedding and leukocyte-endothelial adhesion in postcapillary venules by suppression of matrixmetalloprotease activity with doxycycline. Microcirculation 2009; 16: 657–666.

53. Romanic AM, Harrison SM, Bao W, et al. Myocardial protection from ischemia/reperfusion injury by targeted deletion of matrix metalloproteinase-9. Cardiovasc Res 2002; 54: 549–558.

54. Lukasz A, Hillgruber C, Oberleithner H, et al. Endothelial glycocalyx breakdown is mediated by angiopoietin-2. Cardiovasc Res 2017; 113: 671–680.

55. Drost CC, Rovas A, Kusche-Vihrog K, et al. Tie2 Activation Promotes Protection and Reconstitution of the Endothelial Glycocalyx in Human Sepsis. Thromb Haemost 2019; 119: 1827–1838.

56. Everhardt Queen A, Moerdyk-Schauwecker M, McKee LM, et al. Differential Expression of Inflammatory Cytokines and Stress Genes in Male and Female Mice in Response to a Lipopolysaccharide Challenge. PLoS One 2016; 11: e0152289.

