## Supplementary methods, figures, table for "Matrix Metalloproteinase (MMP) inhibition rescues endothelial glycocalyx damage and reduces neutrophil infiltration in sepsis-associated acute kidney injury"

The inclusion and exclusion criteria are found below

*Sepsis-AKI*

*Inclusion criteria:*

1. Adults over 18 years
2. Sepsis-AKI: suspected sepsis causing organ dysfunction ($\geq$2 SOFA) and evidence of KDIGO AKI stages 1-3
3. Ability to provide informed consent

*Exclusion criteria:*

1. Pregnancy
2. Diabetes
3. Acute stroke
4. Trauma or burns
5. Active cancer

*Healthy Controls*

*Inclusion criteria:*

1. Adults over 18 years

*Exclusion criteria:*

1. Pregnancy
2. Diabetes
3. Acute stroke
4. Trauma or burns
5. Active cancer

### Supplementary Figures

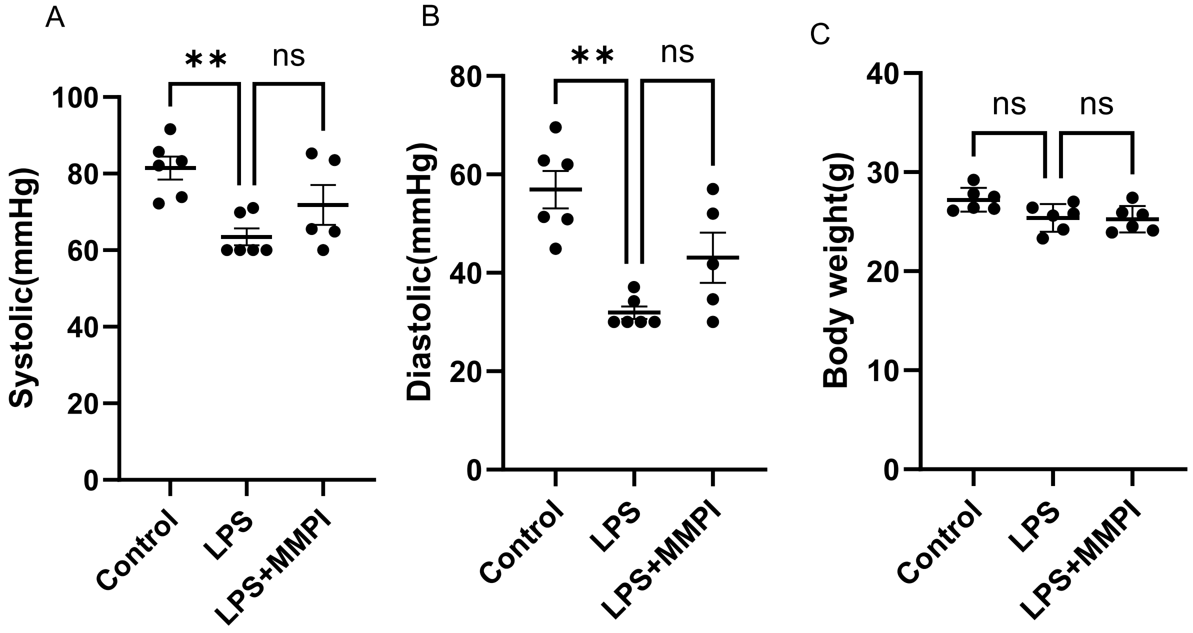


**Supplementary Figure 1. Sepsis-AKI was associated with reduced blood pressure (BP) but no change in body weight.** Systolic and diastolic BP were measured using a noninvasive specialised tail cuff. In the LPS group, 4 mice had systolic and diastolic BP below the detectable range and therefore, the lowest detectable blood pressure was recorded for these mice. Body weight in control, LPS and the LPS+MMPI groups. Each dot on the graph represents a mouse. Data are expressed as the mean ± SEM. One-way ANOVA with Bonferroni's multiple comparison test or, if data are not normally distributed, the Kruskal-Wallis test multiple comparison test was used for statistical analysis.

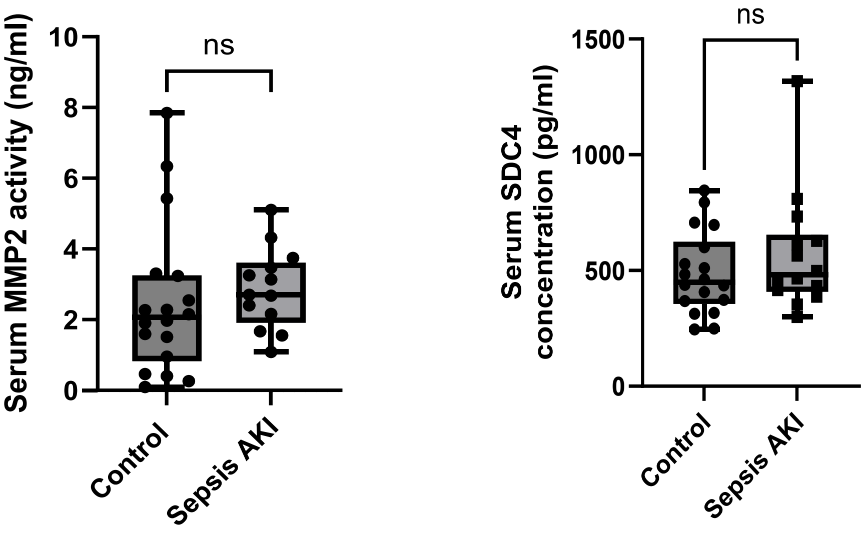


**Supplementary Figure 2: No change in circulating MMP2 activity and SDC4 shedding in sepsis-AKI patients.** Data are expressed as box plots showing median [IQR]. Unpaired t-test was used for statistical analysis. Student t test or Mann-Whitney test was used, dependent on whether the data was normally distributed.

| Genes | Assay ID | Exon Boundary | Assay Location | Amplicon Length |
|---|---|---|---|---|
| SDC1 | Mm00448918_m1 | 2-3 | 387 | 131 |
| SDC4 | Mm00488527_m1 | 4-5 | 469 | 55 |
| ICAM1 | Mm00516023_m1 | 2-3 | 403 | 58 |
| VCAM1 | Mm01320970_m1 | 2-3 | 656 | 71 |
| MMP9 | Mm00442991_m1 | 12-13 | 2098 | 76 |
| MMP2 | Mm00439498_m1 | 2-3 | 674 | 62 |
| GAPDH | Mm99999915_g1 | 2-3 | 117 | 107 |

**Supplementary Table 1** illustrates the list of TaqMan primer probes used in this study
